# Cell Surface β-Lactamase Recruitment: A Facile Selection to Identify Protein-Protein Interactions

**DOI:** 10.1101/2023.07.23.550224

**Authors:** Jordan A. Hinmon, Jade M. King, Latrina J. Mayo, Cierra R. Faries, Ya’hnis T. Street, David W. Crawford, Patrick C. Beardslee, Alexander Hendricks, Brian R. McNaughton

## Abstract

Protein-protein interactions are central to many cellular processes, and the identification of novel protein-protein interactions is a critical step in the discovery of protein therapeutics. Simple methods to identify naturally existing or laboratory evolved protein-protein interactions are therefore valuable research tools. We have developed a facile selection that links protein-protein interaction-dependent β-lactamase recruitment on the surface of E. coli with resistance to ampicillin. Bacteria displaying a protein which form a complex with a specific protein-β-lactamase fusion are protected from ampicillin-dependent cell death. In contrast, bacteria that do not recruit β-lactamase to the cell surface are killed by ampicillin. Given its simplicity and tunability, we anticipate this selection will be a valuable addition to the palette of methods for illuminating and interrogating protein-protein interactions.

## Introduction

The identification and interrogation of protein-protein interactions (PPIs) is necessary to illuminate and better understand complex and disease-relevant cellular processes, as well as to identify novel proteins with therapeutic potential. Facile methods to identify and interrogate PPIs are therefore of great value to a diverse set of researchers.

Popular screening-based methods for detecting and interrogating PPIs include chemical cross-linking^[1]^, co-immunoprecipitation^[2]^, Enzyme-Linked Immunosorbant Assay (ELISA)^[3]^, phage display^[4]^, bacterial display^[5]^, yeast display^[6]^, mRNA display^[7]^, protein-fragment complementation^[8]^, and quantitative proteomic techniques^[9]^. These screening-based methods can be laborious and time consuming, often require specialized and expensive equipment and/or reagents, and can rely on complex data analysis.

In contrast to screens, selection-based methods take advantage of Darwinian evolution, since cells housing partners in a binding interaction survive, while those housing non-binders die. This binary outcome dramatically simplifies the process of identifying interacting partners. Popular selection-based platforms to identify PPIs include bacterial two-hybrid^[10]^, yeast two-hybrid^[11]^, and Phage-Assisted Continuous Evolution (PACE)^[12]^. In each of these methods, a “bait” protein or peptide is fused to a DNA binding domain and a “prey” peptide or protein is fused to an activation domain. Formation of a complex between bait and prey localizes the activation domain into proximity with the promoter of a selection gene necessary for cell survival. While incredibly powerful, each of these methods requires an interaction that orients the activation domain in a position that permits transcription of the selection gene, which can be challenging to predict a priori. While display-based methods allow researchers to alter the concentration of exogenous target (e.g. high prey concentrations may be used in early rounds of screening to identify early-stage hits), tightly controlling the level of the target component using in vivo methods like two-hybrid and PACE can be challenging.

We set out to develop a facile and broadly applicable selection-based display platform to identify two-component PPIs. In our approach, we drew inspiration from bacterial transformation. Selection of bacteria that have been transformed with a plasmid is typically achieved by treatment with an antibiotic, since the plasmid transformed into the bacteria often encodes a gene that endows resistance to a specific antibiotic. For example, many commonly used plasmids contain a gene encoding β-lactamase, which reacts with β-lactam antibiotics (e.g. ampicillin), rendering them non-lethal. We envisaged a selection in which one protein (bait) is displayed on the surface of E. coli, and another protein (prey) is fused to β-lactamase. We reasoned that an interaction between the bait and prey biopolymers would generate a high effective concentration of β-lactamase on the surface of the bacteria, which would provide resistance to a β-lactam antibiotic. In contrast, E. coli lacking a displayed bait with high affinity for the prey would not recruit β-lactamase to the cell surface and would therefore be killed in the presence of a β-lactam antibiotic (**Figure 1a**).

**Figure 1.**
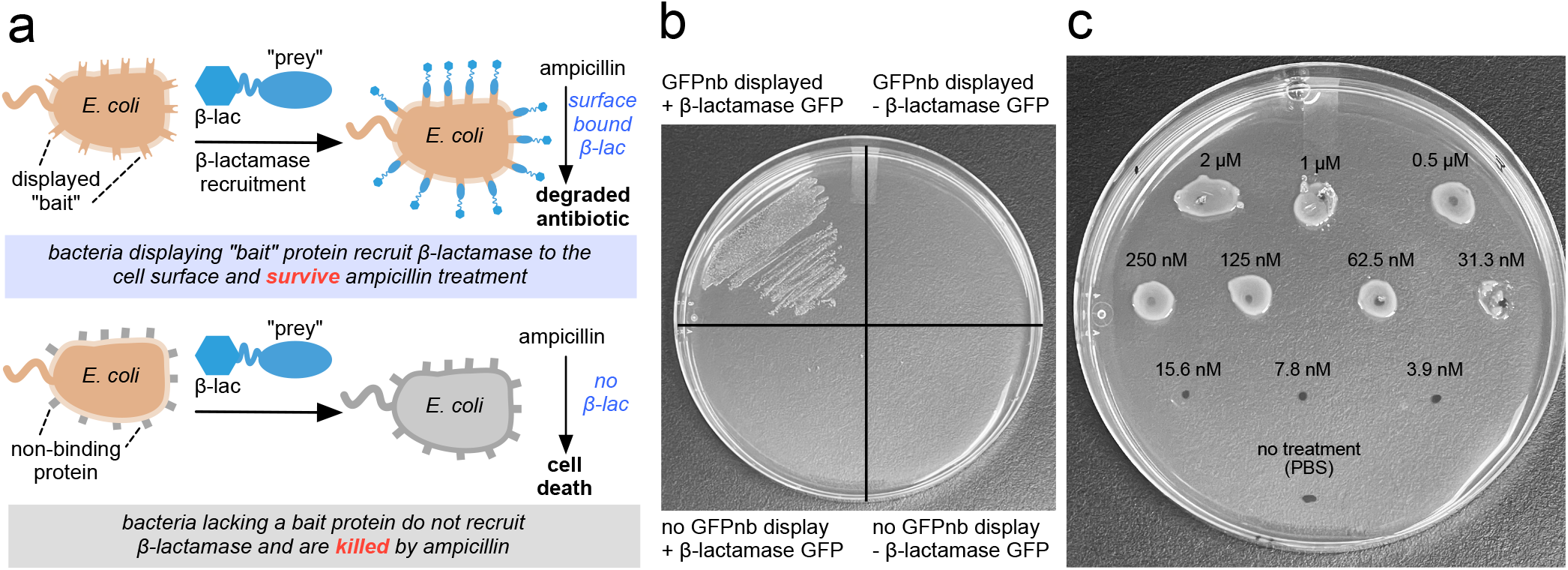
(**a**) Overview of bacterial display β-lactamase recruitment to identify protein-protein interactions. E. coli displaying a bait protein is mixed with a fusion protein consisting of prey, which binds the bait, and β-lactamase. Bait:prey binding generates a high effective concentration of surface bound β-lactamase, which provides resistance to β-lactam antibiotics (e.g. ampicillin). No binding interaction between bait and prey results in cell death in the presence of ampicillin. (**b**) A protein-protein interaction between surface displayed GFP-binding nanobody (GFPnb) and β-lactamase GFP endows resistance to ampicillin (upper left quadrant). E. coli displaying GFPnb that is not incubated with β-lactamase GFP are killed by ampicillin (upper right quadrant). E. coli not displaying GFPnb, but incubated with β-lactamase GFP, is killed with ampicillin (lower left quadrant). E. coli not displaying GFPnb and not treated with β-lactamase GFP is killed with ampicillin (lower right quadrant). (**c**) Survival of E. coli displaying GFPnb is dependent upon the concentration of exogenous β-lactamase GFP used in the bait / prey mixing step of the selection. Selections were performed in triplicate; representative data are shown.

## Results and Discussion

### Protein Interaction-Dependent Cell Surface β-lactamase Recruitment Endows Resistance to Ampicillin

As a proof-of-concept, we transformed E. coli (DH10B-T1R) with pNeae2-GFPnb, a plasmid that endows resistance to chroramphenicol and permits inducible expression of a bacterial display platform (intimin N-terminal domain, Neae^[13]^) fused to a previously reported Green Fluorescent Protein (GFP) binding nanobody^[14]^ (GFPnb, KD ∼1.4 nM) equipped with a C-terminal myc tag. When E. coli was induced with IPTG to display GFPnb-myc, grown for 18 hours at 25 °C, then treated with either AlexaFluor-488 labeled anti-myc antibody or GFP-β-lactamase, we observed high levels of cell surface fluorescence. In contrast, E. coli treated identically, but not induced with IPTG, were not appreciably fluorescent (Supporting Information). Collectively, these data demonstrate that GFPnb display is tightly controlled, when induced with IPTG, GFPnb is displayed at appreciable levels on the cell surface, displayed GFPnb retains affinity for GFP. and GFP recruitment to the exterior of bacteria requires GFPnb display.

We next determined if β-lactamase recruitment provided protection against treatment with β-lactam antibiotic. A 2 mL culture of E. coli (OD600 ∼ 0.4) containing pNeae2-GFPnb was induced with 0.1 mM IPTG to display GFPnb, while another 2 mL culture of identical E. coli was not induced to display GFPnb. After growth for 18 hours at 25 °C. Both induced and uninduced cultures grew to OD600 of approximately 2.5 after overnight growth. From these samples, 200 μL of each was transferred to Eppendorf tubes (two +IPTG samples; two -ITPG samples) and pelleted. 200 μL of a PBS containing 2μMβ-lactamase GFP fusion protein was added to one of the tubes containing E. coli induced to display GFPnb, as well as one of the tubes containing E. coli that was not induced to display GFPnb. The other two tubes were treated with 200 μL PBS. All four tubes were rotated at 25 °C for 30 minutes, pelleted and washed with 500 μL PBS for one minute. After a final pelleting step, E. coli was resuspended in 200 μL PBS and 10 μL of each sample was plated onto LB-agar containing 25 μg/mL chloramphenicol and 100 μg/mL ampicillin.

As shown in **Figure 1b**, following incubation at 30 °C for 20 hours, we observed robust growth of E. coli that was induced to display GFPnb and treated with a solution containing 2μM β-lactamase GFP. In contrast, no growth was observed for the sample induced to display GFPnb but not treated with β-lactamase GFP. Similarly, bacteria not induced to display GFPnb, but treated with 2μM β-lactamase GFP, did not survive, despite treatment with a high concentration of β-lactamase GFP prior to plating. As expected, bacteria not induced to display GFPnb and not treated with β-lactamase GFP did not survive. Collectively, these data demonstrate the viability of the selection platform: cells that recruit β-lactamase via a cell surface PPI survive treatment with ampicillin, while bacteria lacking cell surface recruited β-lactamase are killed by ampicillin.

We next evaluated the concentration-dependence of exogenous β-lactamase GFP on cell survival. Samples of E. coli induced to display GFPnb were individually incubated with 2000 to 4 nM β-lactamase GFP in PBS for 30 minutes at 25 °C. These cells were then pelleted, washed, and resuspended as described above. Next, 5 μL of each resuspended E. coli sample was plated on LB agar containing chloramphenicol and ampicillin. As shown in **Figure 1c**, E. coli treated with 2000 to 31 nM β-lactamase GFP prior to plat-ing survived the selection, while bacteria treated with lower concentrations of β-lactamase GFP did not survive.

### Bacteria Displaying GFPnb are Viable in the β-lactamase Recruitment Selection for at Least Ten Days after Induction

E. coli displaying GFPnb are viable reagents in the selection for at least ten days after IPTG induction. E. coli (OD ∼ 0.4) was induced to display GFPnb with 0.1 mM IPTG, grown at 25 °C for 18 hours, then stored in LB (with IPTG and chloramphenicol) at 4 °C. Over a series of 2, 4, 6, 8, or 10 days after IPTG induction, 200 μL aliquots of E. coli induced to display GFPnb were subjected to selection conditions using 1 μM exogenous β-lactamase GFP. After each selection, cell survival was assessed by plating cells on LB agar containing chloramphenicol and ampicillin. Satisfyingly, as shown in **Figure 2**, we observed simi-lar levels of growth on selection plates, indicating that bacteria displaying GFPnb are viable reagents in the selection, even ten days after IPTG induction. In contrast, identical bacteria that were not induced with IPTG failed to survive the selection (data not shown).

**Figure 2.**
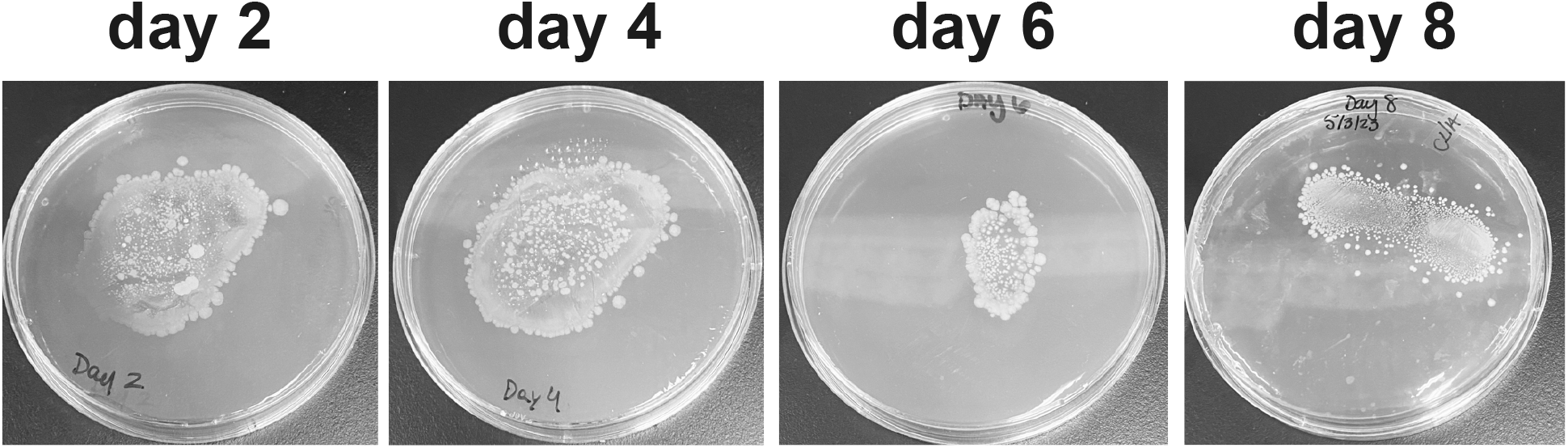
E. coli displaying Neae-GFPnb were treated with 1 μM β-lactamase GFP, washed, then plated on chloramphenicol/ampicillin LB agar 2, 4, 6, or 8 days after IPTG induction. Selections were performed in triplicate; representative data are shown.

### Time-Dependence of the Selection Following IPTG Induction

Having successfully demonstrated the β-lactamase recruitment selection works as envisaged, we de-termined how quickly β-lactamase GFP recruitment and cell survival is achieved following IPTG induction of Neae-GFPnb expression. E. coli harboring pNeae2-GFPnb (OD ∼ 0.4) were induced with 0.1 mM IPTG, then mixed with 1 μM GFP-β-lactamase 1 to 9 hours after induction. Interestingly, appreciable growth, indicating GFPnb display and β-lactamase GFP recruitment, was achieved after 5 hours, and in-creased cell growth was observed for each subsequent time point (**Figure 3a**). In contrast, bacteria not induced with IPTG did not survive on selection plates, over the course of the entire nine-hour experiment (**Figure 3b**). The modest time needed to achieve PPI-dependent cell survival further simplifies the selection protocol.

**Figure 3.**
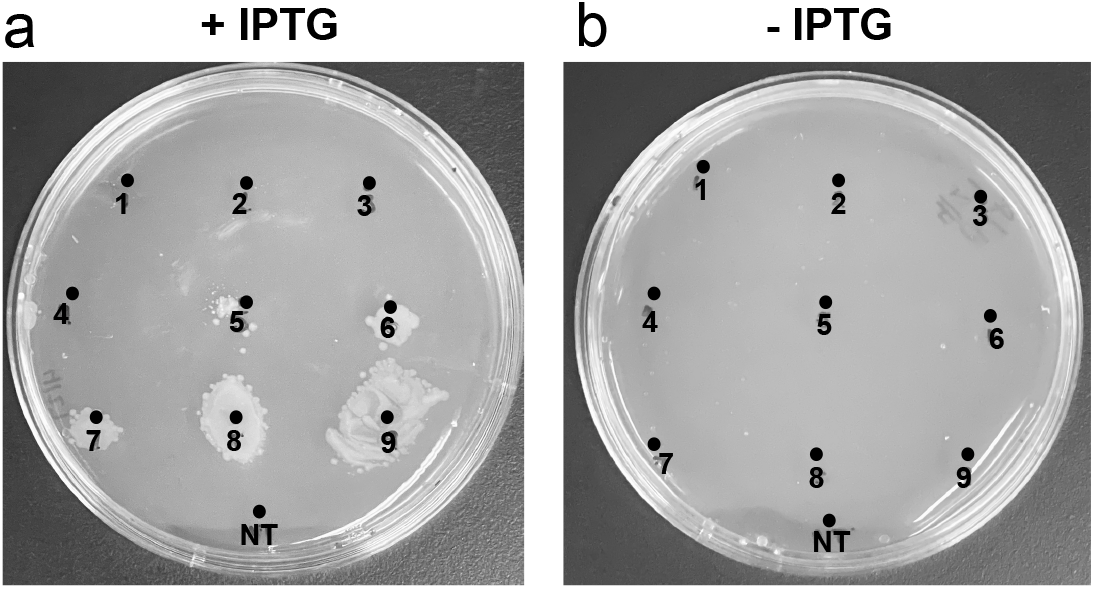
(**a**) E. coli induced to display Neae-GFPnb were were subjected to selection conditions (1 μM β-lactamase GFP and washing) 1-9 hours after IPTG induction, then plated on chloramphenicol/ampicillin LB agar. Experiments were performed in triplicate; representative data are shown. (**b**) E. coli not induced to display Neae-GFPnb were subjected to selection conditions (1 μM β-lactamase GFP and washing) over the identical time-period, then plated on chloramphenicol/ampicillin LB agar. NT = no treatment; cells were not treated with β-lactamase GFP. Experiments were performed in triplicate; representative data are shown.

## Conclusion

By coupling bacterial display of a bait protein with recruitment of an exogenous fusion protein consisting of prey and β-lactamase, we have developed a novel selection to identify PPIs. The selection requires a PPI to recruit a β-lactamase fusion protein to the surface of E. coli, thereby generating a high effective molarity of this enzyme. When β-lactamase fusion protein is recruited to the cell surface, cells survive on LB agar containing ampicillin. In contrast, E. coli lacking recruited β-lactamase fusion protein do not survive in the presence of ampicillin. This selection gives researchers the ability to tightly control the concentration of exogenous prey β-lactamase fusion protein, which should allow researchers to select for high, medium, or low affinity PPIs. Cell survival, resulting from of a desired PPI, is achieved in as little as five hours after IPTG induction of Neae-bait fusion protein, and bacteria displaying bait are viable reagents at least ten days after the initial display of GFPnb was induced. Given the simplicity and tunability of this selection, we anticipate it will be a valuable addition to the palette of experiments commonly used to detect and evaluate PPIs.

## Experimental Section

### Expression and purification of β-lactamase-GFP fusion protein

β-lactamase-GFP was cloned into pET 21a(+), transformed into SHuffle T7 E. coli by heat shock, and the resulting bacteria was plated onto LB-agar containing 100 μg/mL ampicillin. A single colony was selected for future use and prepared as a glycerol stock. From the glycerol stock was grown 5 mL of starter culture in LB containing 100 μg/mL ampicillin (37 °C, 250 RPM, 18 hours). The resulting culture was then added to 250 mL of LB with 100 μg/mL ampicillin, and grown at 37 °C, 250 RPM, to an OD600 ∼0.4. The culture was removed from the incubator, allowed to cool to room temperature, then induced with IPTG (0.4 mM final concentration). Bacteria was then grown overnight at 25 °C, 250 RPM for approximately 18 hours. Bacteria was pelleted by centrifugation, lysed with complete B-Per, and purified by Ni-NTA chromatography. Following washing steps with 10 mM, then 25 mM histidine, protein was eluted with 250 mM histidine. Protein purity was characterized by SDS-PAGE (4-10% Tricine precast gel) and stained with InstantBlue Coomassie.

### Display of Neae-GFPnb fusion proteins

pNeae2-GFPnb-myc was transformed into DH10B-T1R E. coli by heat shock, and the resulting bacteria was plated onto LB-agar containing 25 μg/mL chloramphenicol. A single colony was selected for future use and prepared as a glycerol stock. Bacteria was grown as a small (2-5 mL) culture in LB with 25 μg/mL chloramphenicol at 37 °C, shaking at 250 RPM, until an OD600 ∼0.4 was reached. E. coli was then induced with IPTG (final concentration = 0.1 mM) and grown at 25 °C for approximately 12-18 hours.

### Selection protocol

DH10B-T1R E. coli transformed with pNeae-GFPnb was grown in 2 mL LB containing 25 μg/mL chloramphenicol at 37 °C, shaking at 250 RPM, until an OD600 ∼0.4 was reached. E. coli was then induced with IPTG (final concentration = 0.1 mM) and grown at 25 °C (250 RPM) for approximately 12 hours. 200 μL of bacteria was transferred to a 2 mL Eppendorf tube and pelleted at 10,000 RPM for 2 minutes. Supernatant was disposed and cells were resuspended in 200 μL of PBS containing 2 μM β-lactamase-GFP. Cells were mixed on a shaker (40 RPM) for 30 minutes at room temperature, pelleted, and resuspended in 500 μL of PBS. Cells were then mixed by rotation for 1 minute at room temperature. Following this washing step, bacteria was pelleted and resuspended in 200 μL of PBS. 10 μL of this solution was spread on LB agar plates containing 25 μg/mL chloramphenicol and 100 μg/mL ampicillin and placed in a 30 °C incubator for 18 hours.

## Acknowledgements

Delaware State University is acknowledged for institutional support. We thank Michael Moore (DSU) from the Optical Science Center for Applied Research (OSCAR) imaging facility for assisting with fluorescence imaging experiments. This work was supported by the National Institutes of Health/National Institute of General Medical Sciences (R15GM141783, B.R.M.) and National Institutes of Health/National Institute on Minority Health and Health Disparities (U54MD015959, B.R.M.). C.R.F. was funded by a grant from the National Institutes of Health/National Institute of General Medical Sciences (T34GM136477).

## Conflict of Interests

The authors declare no conflict of interest.

## Notes

### Competing Interest Statement

The authors have declared no competing interest.

