## supporting information and figures for "Cell Surface β-Lactamase Recruitment: A Facile Selection to Identify Protein-Protein Interactions"

#### Materials

Phosphate-Buffered Saline, 1X, Corning
Alexa Fluor 488 Anti-Myc tag antibody, Abcam (ab202008)
LB Miller Broth, Fisher
Agar, Fisher
Ampicillin, GoldBio Technology
Chloramphenicol, Thermo Scientific
Isopropyl-b-D-1-thiogalactopyranoside (IPTG), GoldBio Technology
HisPur Ni-NTA Resin, Thermo Scientific
PageRuler Prestained Protein Ladder, Thermo Scientific
4-20% Tricine precast gels, Thermo Scientific
SHuffle T7 Express Competent *E. coli*, NEB
Complete B-Per cell lysis reagent, Thermo Scientific
2 mL Eppendorf tubes, Thermo Scientific
Culture tubes, Thermo Scientific
InstantBlue Coomassie, Thermo Scientific

#### Abbreviations

GFP: Green Fluorescent Protein
GFPnb: GFP nanobody
LB: Luria Broth
PBS: phosphate buffered saline
*E. coli*: *Escherichia coli*
AF488: Alexafluor-488
RPM: rotations per minute
$OD_{600}$: optical density, $\lambda = 600$ nm

#### Instrumentation and Software

Shaking Incubator: MaxQ, Thermo Scientific
Fluorescence microscopy: Zeiss 780 confocal with spectral detection. 63X 1.4 NA Plan Apochromat oil objective
Protein concentration: NanoDrop One was used to measure $Abs_{280}$. Protein extinction coefficient was calculated using https://www.aatbio.com/tools/calculate-protein-concentration and https://www.bioline.com/media/calculator/01_04.html
*E. coli* concentration: $OD_{600}$ was measured using a NanoDrop One
Sequence logos: WebLogo, https://weblogo.berkeley.edu/logo.cgi
Sequence analysis: Geneious

#### Protein sequences

All plasmids were ordered from Genescript

##### Neae-(E tag)-GFPnb-myc

MITHGCYTRTRHKHKLKKTLIMLSAGLGLFFYVNQNSFANGENYFKLGSDSKLLTHDSYQNRLFYTLKTG
ETVADLSKSQDINLSTIWSLNKHLYSSESEMMKAAPGQQIILPLKKLPFEYSALPLLGSAPLVAAGGVAG
HTNKLTKMSPDVTKSNMTDDKALNYAAQQAASLGSQLQSRSLNGDYAKDTALGIAGNQASSQLQAWLQHY
GTAEVNLQSGNNFDGSSLDFLLPFYDSEKMLAFGQVGARYIDSRFTANLGAGQRFFLPANMLGYNVFIDQ
DFSGDNTRLGIGGEYWRDYFKSSVNGYFRMSGWHESYNKKDYDERPANGFDIRFNGYLPSYPALGAKLIY
EQYYGDNVALFNSDKLQSNPGAATVGVNYTPIPLVTMGIDYRHGTGNENDLLYSMQFRYQFDKSWSQQIE
PQYVNELRTLSGSRYDLVQRNNNIILEYKKQDILSLNIPHDINGTEHSTQKIQLIVKSKYGLDRIVWDDS
ALRSQGGQIQHSGSQSAQDYQAILPAYVQGGSNIYKVTARAYDRNGNSSNNVQLTITVLSNGQVVDQVGV
TDFTADKTSAKADNADTITYTATVKKNGVAQANVPVSFNIVSGTATLGANSAKTDANGKATVTLKSSTPG
QVVVSAKTAEMTSALNASAVIFFDGAPVPYPDPLEPAQPAMQVQLVESGGALVQPGGSLRLSCAASGFPV
NRYSMRWYRQAPGKEREWVAGMSSAGDRSSYEDSVKGRFTISRDDARNTVYLQMNSLKPEDTAVYYCNVN
VGFEYWGQGTQVTVSSAAAEQKLISEEDLAA***

##### His6-β-lactamase-GFP

MGSSHHHHHHSQDPHPETLVKVKDAEDQLGARVGYIELDLNSGKILESFRPEERFPMMSTFKVLLCGAVL
SRIDAGQEQLGRRIHYSQNDLVEYSPVTEKHLTDGMTVRELCSAAITMSDNTAANLLLTTIGGPKELTAF
LHNMGDHVTRLDRWEPELNEAIPNDERDTTMPVAMATTLRKLLTGELLTLASRQQLIDWMEADKVAGPLL
RSALPAGWFIADKSGAGERGSRGIIAALGPDGKPSRIVVIYTTGSQATMDERNRQIAEIGASLIKHWGGG
GSGGGGSMVSKGEELFTGVVPILVELDGDVNGHKFSVSGEGEGDATYGKLTLKFICTTGKLPVPWPTLVT
TLTYGVQCFSRYPDHMKQHDFFKSAMPEGYVQERTIFFKDDGNYKTRAEVKFEGDTLVNRIELKGIDFKE
DGNILGHKLEYNYNSHNVYIMADKQKNGIKVNFKIRHNIEDGSVQLADHYQQNTPIGDGPVLLPDNHYLS
TQSALSKDPNEKRDHMVLLEFVTAAGITLGMDELYK***

#### Experimental Methods

##### Expression and purification of β-lactamase-GFP fusion protein

β-lactamase-GFP was cloned into pET 21a(+), transformed into SHuffle T7 *E. coli* by heat shock, and the resulting bacteria was plated onto LB-agar containing 100 μg/mL ampicillin. A single colony was selected for future use and prepared as a glycerol stock. From the glycerol stock was grown 5 mL of starter culture in LB containing 100 μg/mL ampicillin (37 °C, 250 RPM, 18 hours). The resulting culture was then added to 250 mL of LB with 100 μg/mL ampicillin, and grown at 37 °C, 250 RPM, to an $OD_{600}$ ~0.4. The culture was removed from the incubator, allowed to cool to room temperature, then induced with IPTG (0.4 mM final concentration). Bacteria was then grown overnight at 25 °C, 250 RPM for approximately 18 hours. Bacteria was pelleted by centrifugation, lysed with complete B-Per, and purified by Ni-NTA chromatography. Following washing steps with 10 mM, then 25 mM histidine, protein was eluted with 250 mM histidine. Protein purity was characterized by SDS-PAGE (4-10% Tricine precast gel) and stained with InstantBlue Coomassie.

##### Display of Neae-GFPnb fusion proteins

pNeae2-GFPnb-myc was transformed into DH10B-$T1^R$ *E. coli* by heat shock, and the resulting bacteria was plated onto LB-agar containing 25 μg/mL chloramphenicol. A single colony was selected for future use and prepared as a glycerol stock. Bacteria was grown as a small (2-5 mL) culture in LB with 25 μg/mL chloramphenicol at 37 °C, shaking at 250 RPM, until an $OD_{600}$ ~0.4 was reached. *E. coli* was then induced with IPTG (final concentration = 0.1 mM) and grown at 25 °C for approximately 12-18 hours.

##### Fluorescence microscopy analysis of nanobody and β-lactamase-GFP recruitment

Duplicates of DH10B-$T1^R$ *E. coli* previously transformed with pNeae2-GFPnb-myc or pNeae2-GFPnb(off)-myc were grown in 2 mL LB containing 25 μg/mL chloramphenicol at 37 °C, shaking at 250 RPM, until an $OD_{600}$ ~0.4 was reached. One culture of DH10B-$T1^R$ *E. coli* transformed with pNeae2-GFPnb-myc, or pNeae2-GFPnb(off)-myc, were then treated with IPTG (final concentration = 0.1 mM). The other two cultures of DH10B-$T1^R$ transformed with pNeae2-GFPnb-myc, or pNeae2-GFPnb(off)-myc were not induced. All four samples (pNeae2-GFPnb-myc / +IPTG; pNeae2-GFPnb-myc / -IPTG; pNeae2-GFPnb(off)-myc / +IPTG; pNeae2-GFPnb(off)-myc / -IPTG) were grown at 25 °C for approximately 18 hours. Duplicates of each culture (200 µL) were transferred to eight separate 2 mL Eppendorf tubes and pelleted at 10,000 RPM for 2 minutes. Supernatant was disposed and cells were resuspended in 200 µL of PBS containing either 2 µM β-lactamase-GFP, or 200 µL of PBS with 2 µL anti-*myc* AF488 antibody. Cells were mixed on a shaker (40 RPM) for 30 minutes at room temperature, pelleted, and resuspended in 500 µL of PBS. Cells were then mixed by rotation for 1 minute at room temperature. Following this washing step, bacteria were pelleted and resuspended in 200 µL of PBS. 10 µL of each solution was pipetted onto a glass slide, covered with a glass cover slip, and placed in aluminum foil until fluorescence imaging. Imaging was done using a Zeiss 780 confocal with spectral detection. 63X 1.4 NA Plan Apochromat oil objective. Excited with 488nm laser and imaged using the spectral detector with a detection band of 505-560.

### Supporting Information

**Selection protocol**
DH10B-T1$^R$ *E. coli* transformed with pNeae-GFPnb was grown in 2 mL LB containing 25 μg/mL chloramphenicol at 37 °C, shaking at 250 RPM, until an $OD_{600}$ ~0.4 was reached. *E. coli* was then induced with IPTG (final concentration = 0.1 mM) and grown at 25 °C (250 RPM) for approximately 12 hours. 200 μL of bacteria was transferred to a 2 mL Eppendorf tube and pelleted at 10,000 RPM for 2 minutes. Supernatant was disposed and cells were resuspended in 200 μL of PBS containing 2 μM β-lactamase-GFP. Cells were mixed on a shaker (40 RPM) for 30 minutes at room temperature, pelleted, and resuspended in 500 μL of PBS. Cells were then mixed by rotation for 1 minute at room temperature. Following this washing step, bacteria was pelleted and resuspended in 200 μL of PBS. 10 μL of this solution was spread on LB agar plates containing 25 μg/mL chloramphenicol and 100 μg/mL ampicillin and placed in a 30 °C incubator for 18 hours.

**β-lactamase-GFP concentration-dependence on cell survival**
DH10B-T1$^R$ *E. coli* containing pNeae-GFPnb were grown in LB with 25 μg/mL chloramphenicol at 37 °C, shaking at 250 RPM, until an $OD_{600}$ ~0.4 was reached. *E. coli* was then induced with IPTG (final concentration = 0.1 mM) and grown at 25 °C (250 RPM) for approximately 12 hours. 200 μL of bacteria were transferred to a 2 mL Eppendorf tube and pelleted at 10,000 RPM for 2 minutes. Supernatant was disposed and cells were resuspended in 200 μL of PBS containing 2 μM to 3.9 nM β-lactamase-GFP. Cells were rotated (40 RPM) for 30 minutes at room temperature, pelleted and resuspended in 500 μL of PBS and mixed by rotation for 1 minute at room temperature. Following this washing step, bacteria was pelleted and resuspended in 200 μL of PBS. 10 μL of this solution was applied to LB agar plates containing 25 μg/mL chloramphenicol and 100 μg/mL of ampicillin. Bacteria solutions were spread into small circles with a pipette tip and the plates were placed in a 30 °C incubator overnight for 18 hours.

**Time-dependence of IPTG induction on cell survival**
DH10B-T1$^R$ *E. coli* containing pNeae-GFPnb were grown in LB with 25 μg/mL chloramphenicol at 37 °C, shaking at 250 RPM, until an $OD_{600}$ ~0.4 was reached. *E. coli* was then induced with IPTG (final concentration = 0.1 mM) and grown at 25 °C (250 RPM) for 0.25 - 8 hours. After each tested timepoint, 200 μL of bacteria were transferred to a 2 mL Eppendorf tube and pelleted at 10,000 RPM for 2 minutes. Supernatant was disposed and cells were resuspended in 200 μL of PBS containing 1 μM β-lactamase-GFP. Cells were rotated (40 RPM) for 20 minutes at room temperature, pelleted and resuspended in 500 μL of PBS and mixed by rotation for 1 minute at room temperature. Following this washing step, bacteria was pelleted and resuspended in 200 μL of PBS. 10 μL of this solution was applied to LB agar plates containing 25 μg/mL chloramphenicol and 100 μg/mL of ampicillin. Bacteria solutions were spread into small circles with a pipette tip and the plates were placed in a 30 °C incubator overnight for 18 hours.

#### Supporting Figures

**Figure S1. SDS-PAGE of purified β-lactamase-GFP (MW ≈ 58 kDa)**
*The left lane is PageRuler prestained protein ladder, and the right lane is purified β-lactamase-GFP. SDS-PAGE was performed using a Tricine 4-10% precast gel, which was stained with InstantBlue Coomassie.*

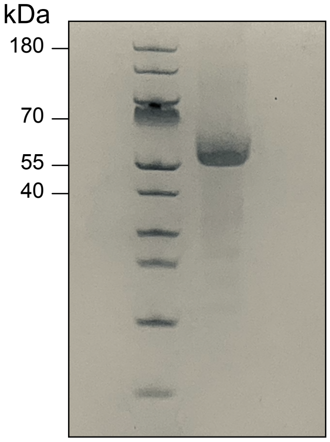

**Figure S2. Fluorescence microscopy images of bacteria containing pNeae2-GFPnb-myc and following treatment with β-lactamase-GFP or anti-myc alexafluor-488 +/- IPTG induction**

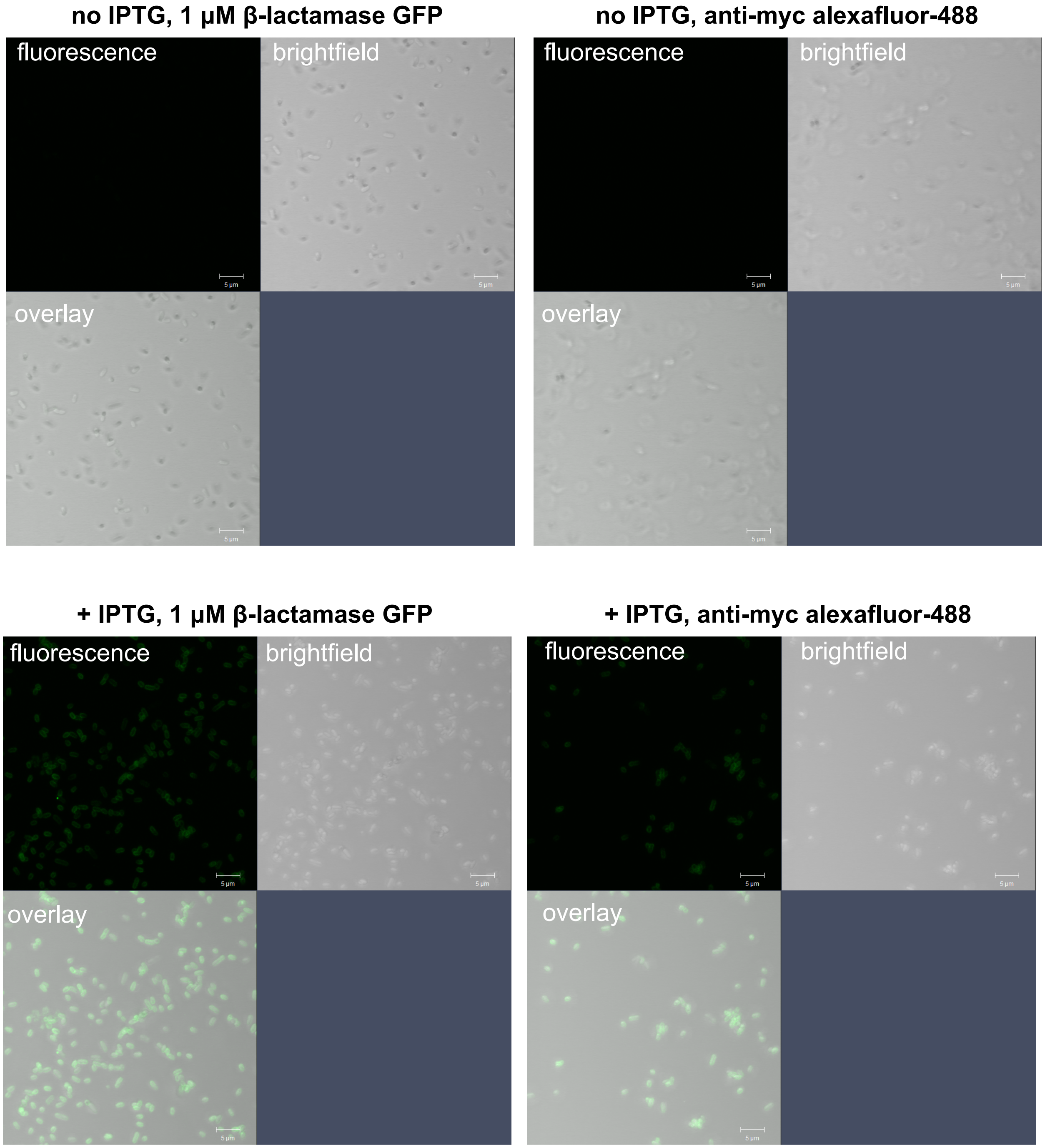
